# In vivo evaluation of a novel compliance-matching vascular graft

**DOI:** 10.1101/2023.11.10.566623

**Authors:** G. Rovas, P. Reymond, M. van Steenberghe, J. Diaper, V. Bikia, M. Cikirikcioglu, W. Habre, C. Huber, N. Stergiopulos

## Abstract

**Background:** The mismatch of elastic properties between the arterial tissue and the vascular grafts, commonly called compliance mismatch, is responsible for many deleterious post-operative complications. Currently, there is an absence of prostheses that conform with the compliance of healthy aortas.

**Objectives:** We proposed a novel compliance-matching graft design, composed of a standard aortic graft surrounded by an optimized Nickel-Titanium compliance-augmenting layer. We aimed to evaluate the in vivo performance of the novel grafts in a swine model and compare it to the native aorta and to gold-standard aortic grafts.

**Methods:** We replaced the thoracic aorta of six domestic pigs with compliance-matching grafts under cardiopulmonary bypass. We removed the compliance-regulating layer of the compliant grafts, so that gold-standard grafts remained implanted.

**Results:** The compliance-matching grafts were implanted without surgical complications and without inducing post-operative hypertension by maintaining systolic pressure (11% increase), aortic pulse wave velocity (17% decrease) and aortic distensibility (40% increase) at healthy levels. The gold-standard grafts caused a significant rise in systolic pressure (47%), pulse pressure (126%) and pulse wave velocity (64%).

**Conclusions:** Our novel compliant grafts could diminish the complications caused by compliance-mismatch and they could surpass the clinical performance of existing prostheses. The proposed grafts comprise a step towards optimized treatment and improved life expectancy of patients subjected to aortic replacement.

## 1. Introduction

Vascular grafts have been used in the treatment of various cardiovascular diseases for many years. In spite of their broad adoption in vascular replacement and bypass surgery, and the numerous improvements they received in terms of design and materials, synthetic vascular grafts are still incapable of mimicking the elastomechanical properties of healthy arteries ^1^. The existing grafts remain multiple times stiffer than the native arterial tissue ^2,3^, leading to a phenomenon called compliance mismatch ^4,5^. The first indications of compliance mismatch have been reported shortly after the establishment of vascular grafts by experiments involving aortic bypass grafting in dogs ^6^. Since then, there has been a notable lack of grafts that could resolve this issue, especially in the case of large-diameter grafts, such as those used for aortic replacement. Regardless of size, compliance mismatch in combination with other suboptimal characteristics, such as the deterioration of mechanical properties and dilatation with time, results in the inadequate performance and poor surgical outcomes of existing grafts ^7^. Consequently, current grafts fail to adequately reduce the burden of cardiovascular diseases that still constitute the leading cause of death and disability worldwide, with a continuously rising tendency ^8^.

Although the root of compliance mismatch is common across grafts of different sizes, we can differentiate between grafts of large and small caliber as far as the complications caused by reduced compliance are concerned. Non-compliant large-diameter grafts cause loss of the Windkessel function of large arteries and reduction in the total arterial compliance, since two-thirds of the total arterial compliance resides in the aorta ^9^. The detrimental effects of reduced arterial compliance on the cardiovascular system have been intensively researched because of the role of arterial compliance in cardiovascular aging and in the genesis of essential hypertension ^10–12^. In association with these findings, vascular grafts have been shown to impair heart function by often causing hypertension, left ventricular hypertrophy, decreased coronary flow and myocardial ischemia ^6,13–17^. On the contrary, the mismatched elastic properties of small-caliber grafts are responsible for more localized but equally deleterious effects, including intimal hyperplasia and thrombosis ^4^. In both types of grafts, the abrupt change of mechanical properties at the anastomoses has been associated with additional complications such as rupture, infection, re-stenosis, migration and endoleak ^18–20^.

In the recent literature there is a lack of large-diameter vascular grafts designs that can match the compliance of human aortas, and there are even fewer such grafts that have been evaluated in animal models. The most notable attempt was a biomimetic graft inspired by the body structure of caterpillars with two polymer layers ^21^. This graft achieved higher compliance than existing alternatives, but it failed to cover the entire physiological range of human aortic compliance, while its performance has not been evaluated in vivo. Conversely, a plethora of promising designs for small-diameter grafts have been proposed and assessed in vivo. They range from implementations based on new materials or material combinations ^22–26^ to cellularized biodegradable grafts ^27,28^. These approaches can reach the compliance of peripheral arteries but not the aortic compliance, which is generally several times higher. Small-diameter grafts are, also, unlikely to be effective for large artery replacement due to the difference in design requirements and operating conditions.

Here, we aimed to evaluate acutely the in vivo performance of a novel compliance-matching (CM) aortic graft, which achieved physiological levels of compliance in vitro ^29^. The compliant graft consisted of a gold-standard graft, surrounded by an optimized compliance-regulating outer layer. We hypothesized that the compliant graft would not alter the hemodynamic conditions when compared to the native aorta. To verify this hypothesis, we surgically replaced the thoracic aorta of healthy swine with the compliant graft by open heart surgery under cardiopulmonary bypass. Thus, we were able to measure under the same conditions the acute hemodynamic effects of the compliant grafts on the cardiovascular system and compare them to healthy aortas and commercial grafts.

## 2. Materials and methods

### 2.1 Compliance-matching vascular grafts

We have described the design, optimization and in vitro evaluation processes of the biomimetic, compliance-matching graft in our previous publication ^29^. In brief, we designed a multi-layer vascular graft that can mimic the elastomechanical properties of a healthy human aorta. The graft visually resembled a vascular stent-graft, but there are fundamental differences in terms of function, design and properties between the proposed CM graft and commercial stent-grafts. The CM graft consisted of a standard, woven, Polyethylene Terephthalate (PET/Dacron) vascular graft (Gelweave,Terumo Corporation, Tokyo, Japan) as an inner layer, and a superelastic, Nickel Titanium (Nitinol/NiTi) exoskeleton, which was similar in appearance to a vascular stent (Fig. 1). The purpose of the stent exoskeleton was to radially compress the inner layer and control its expansion, by altering the characteristic pressure-diameter curve of the graft. The compression generated stored length circumferentially, so that the inner layer performed a coiling and uncoiling movement within the physiological pressure variation, while most of the circumferential load was borne by the stents. As a result, the whole graft acquired increased compliance, even though the inner layer was inelastic. We optimized the design parameters of the stents via a validated computer simulation until the compliance of the whole graft reached physiological human levels ^30^. The optimized stents were laser-cut, inserted over the inner layer with a custom mandrel, and secured in place with surgical suture. We sterilized all the components prior to the assembly, and we sterilized the assembly environment and the assembled grafts via UV radiation. The sterilization process did not alter the properties of the graft. Based on a cadaveric study, we selected inner grafts with length of 15 cm and nominal diameter of 16 mm, whereas the optimization process resulted in a compliance-matching graft whose unloaded, internal diameter was 14mm (lower than the inner graft due to compression).

**Figure 1.**
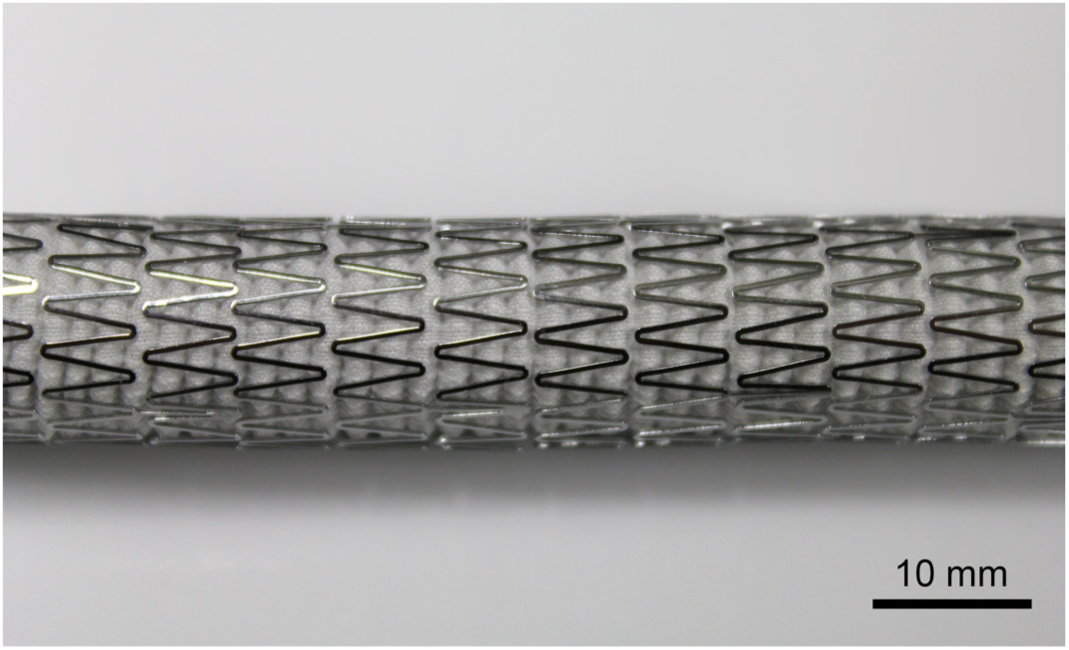
Photograph of the compliance-matching graft, which was used in the experiments.

### 2.2 Surgical procedure

Six, healthy, female, domestic pigs with a mean weight of 49 kg were premedicated by intramuscular injection of a mixture of azaperone (8 mg/kg), midazolam (0.75 mg/kg) and atropine (25 μg/kg). They were intubated under anesthesia with intravenous administration of fentanyl (2 μg/kg) and atracurium (0.5 mg/kg). Anesthesia was maintained with inhalation of sevoflurane (2−3% v/v) and continuous intravenous infusion of fentanyl (10 μg/(kg/h)) and midazolam (0.1 mg/(kg/h)). They were mechanically ventilated with controlled volume mode at a constant insufflation rate (6 ml/kg) using a ventilator with additional software (Servo-i, Maquet Critical Care Ab, Solna, Sweden), at a frequency of 16−20 min^-1^, with the aim of obtaining normal expiratory concentration of CO_2_ (5.5−6 kPa). A positive end-expiratory pressure of 5 cmH_2_O was applied and the fraction of inspired oxygen was set to 0.4 throughout the procedure. Once the anesthesia and the level of analgesia were assured, the right femoral artery was cannulated using a catheter for continuous pressure monitoring. Rectal temperature was monitored using a temperature sensor (Thermalert TH-8, Physitemp, Clifton, NJ) and it was maintained at 38 °C using a heating pad and a heating blanket (Bair Hugger, 3M, Maplewood, MN). We performed a transesophageal echocardiographic examination to assess the myocardial contractility and the aortic diameter.

After ensuring hemodynamic stability, sternotomy was performed followed by hemostasis of the sternum and separation of the two sternal edges with a sternum retractor. Heparin was administered intravenously (300 IU/kg) and the heparinization was repeated, if necessary, to achieve an activated coagulation time (ACT) higher than 400 s. After pericardiotomy, the aorta and the aortic arch were dissected free from the surrounding tissues. The aorta was cannulated using a 16F arterial cannula (Edwards Lifesciences, Irvine, CA) and the right atrium using a single cannula approach.

### 2.3 Aortic pressure and flow rate measurement

We temporarily secured an appropriately-sized, perivascular, ultrasonic flow probe (Transonic Systems Inc., Ithaca, NY) around the ascending aorta, ensuring good acoustic coupling with ultrasound gel. Additionally, we measured the aortic pressure invasively with a calibrated pressure transducer (ADInstruments Inc., Colorado Springs, CO) following standard methodology. The signals from the flow and pressure transducers were recorded simultaneously on a PowerLab data acquisition device (ADInstruments, Dunedin, New Zealand) at a sampling frequency of 1 kHz. After ensuring that the hemodynamic condition was stable, we recorded the two signals until 40 consecutive, artifact-free, cardiac cycles were captured. This recording constituted the after-instrumentation, control measurement [Control]. Then, we removed the two sensors and continued the operation.

### 2.4 Cardiopulmonary bypass and aortic replacement

We initiated the extracorporeal circulation circuit (ECC) consisting of a heart-lung machine (Sorin Stockert SIII, LivaNova, London, United Kingdom) and a membrane oxygenator, following heparinization of the circuit (300 IU/kg). Hypothermia at 28 °C was gradually induced with the heat exchanger. Then, the cardioplegia solution (Cardioplexol + 300mg Procain) was administered once (2 x 50 ml) until the complete cessation of electrical activity. Purse-string sutures were placed on the right auricle and on the aorta, opposite of the first supra-aortic trunk, allowing selective cannulation during circulatory arrest. A 15F and a 27F Bio-Medicus cannulas (Medtronic, Minneapolis, MN) were inserted in the supra-aortic trunk and the right auricle, respectively, and they were tightened with the purse-string sutures. Cerebral perfusion was maintained via the supra-aortic cannula at the theoretical flow rate (8−10 ml/kg). The aortic arch was dissected by a complete transverse incision so that the island of the aortic arch with the two supra-aortic trunks and the perfusion cannula remained united, while the lower part of the aortic arch was excised. This was necessary to achieve selective cerebral perfusion. Subsequently, the compliance-matching graft was sutured distally, at the most distal part of the descending aorta that the anatomy of the thoracic cavity allowed, and so that at least 10 cm of the graft would be implanted. The proximal side of the graft was then sutured at the sinotubular junction. A 15F Bio-Medicus cannula was inserted in the proximal side of the graft and was secured with a purse-string suture. The placement of the cannulas that were necessary for the connection to the ECC can be seen in Fig. 2.

**Figure 2.**
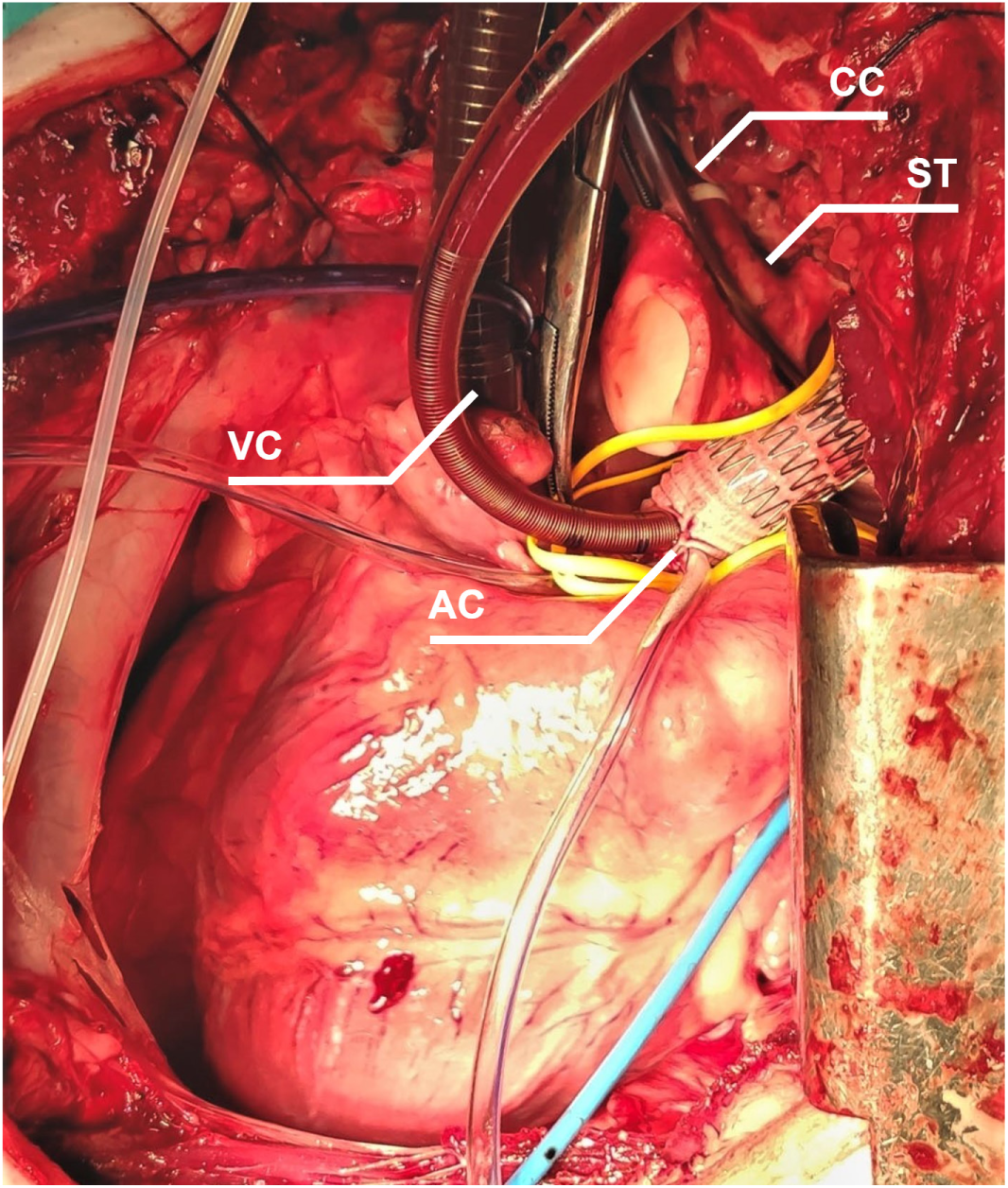
Overview of the surgical procedure and the cannulation utilized for the cardiopulmonary bypass. We performed standard cannulation at the right auricle via the venous cannula (VC) and at the aortic graft via the aortic cannula (AC). Cerebral perfusion was achieved via the cerebral cannula (CC), which was inserted in the island consisting of the aortic arch and the two supra-aortic trunks (ST). In the figure, the aorta has been replaced by the compliance-matching graft. The cannulas and the equipment remained in place after the removal of the stents.

The animal was gradually rewarmed (1°C every 5−10 min) at 36 °C, and the grafted aorta along with the cardiac chambers were purged via a cardioplegia needle. The hemostasis was verified, the ACT was checked every 20−30 min to verify that it remains higher than 400 s, protamine was administered, and the heart was restarted. On some occasions, electric shock was necessary to restore cardiac function. The flow of the ECC was gradually reduced, until the animal was no longer under cardiopulmonary bypass. To reduce the duration and the complexity of the surgery, we decided not to anastomose the island with the supra-aortic trunks on the graft, and we maintained selective cerebral perfusion via the ECC. The cerebral perfusion and oxygenation were monitored with near-infrared spectroscopy (NIRS) throughout the surgery to ensure the preservation of intact autonomic reflex. Additionally, we monitored the arterial blood gas levels along with the ECC blood flow rate, pressure, and temperature until the end of the surgery.

### 2.5 Measurements with the compliant grafts and the PET grafts implanted

The thoracic aorta was replaced by the compliance-matching graft. We waited for the stabilization of the hemodynamic conditions, and we repeated the aortic pressure and blood flow measurements with the same procedure as in 2.3. This measurement constituted the compliance-matching [CM] graft recording.

We secured the pressure and flow sensors in place, and we removed the stents of the compliance-matching graft by cutting them and detaching them from the inner layer. Hence, only the reference, PET graft, which served as the inner layer, remained implanted. We repeated the acquisition of pressure and flow rate, and this final measurement comprised the recording with the gold-standard graft, which is named hereinafter from the material of the reference graft [PET]. The three stages of the surgical procedure during which we performed the intra-operative measurements can be seen in Fig. 3. After the final measurements, the animals were sacrificed while they were still under general anesthesia.

**Figure 3.**
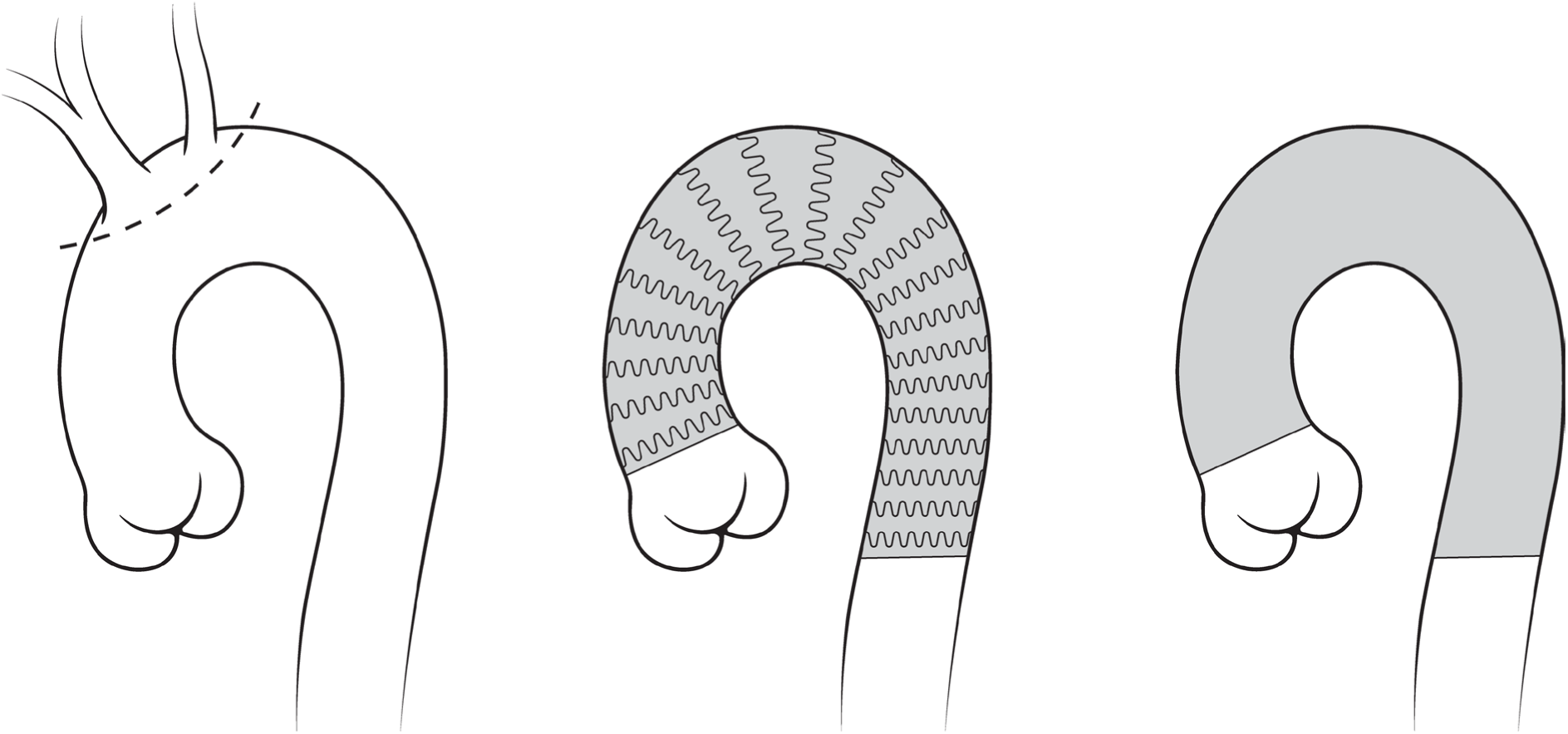
The three stages of the surgical procedure. We recorded aortic pressure and flow rate at three stages during the surgery: with the native aorta [Control], with the compliance-matching graft [CM], and with the gold-standard graft [PET].

The animal experiments protocol was approved by the University of Geneva ethics committee and by the cantonal Consumer and Veterinary Affairs Department [Cantonal no.: GE116, Federal no.: 33865], and it conformed with the all the laws and regulations in force regarding animal welfare and animal experimentation [SR 455, SR 455.1, SR 455.163].

### 2.6 Data analysis

The aortic pressure and flow rate were directly measured by the implanted transducers, and they were processed offline. All subsequent analyses were automatically performed with custom software programmed in Matlab (MathWorks, Natick, Massachusetts) without user interference. We followed the standard methodology to calculate the hemodynamic parameters and to perform pulse-wave analysis. Both signals were filtered with a Savitzky–Golay filter with a window length of 81 ms to remove high frequency noise, and then they were segmented into cardiac cycles. We rejected outlier cardiac cycles whose pressure laid outside the “envelope” defined by the mean ± two standard deviations. We calculated the heart rate and the systolic, diastolic, mean and pulse pressure based on the recorded pressure signal, while from the flow signal we calculated the maximum and minimum flow rate as well as the cardiac output and stroke volume. These parameters were calculated separately over the recorded heartbeats and then averaged to obtain the mean values.

We derived the peripheral resistance *R*_*P*_ as the ratio of mean aortic pressure over mean aortic flow rate and then used it to calculate the total arterial compliance *C*_*T*_ based on the pulse pressure method ^31^. The aortic characteristic impedance *Z*_*C*_ was derived as the mean ratio of pressure over flow rate in the frequency domain, averaged over the third to tenth harmonic, excluding harmonics whose distance from the mean was more than twice the standard deviation ^32^. Before the sternotomy, we measured the diameter of the ascending aorta using transesophageal, M-mode echography, while the diameters of the CM graft and the PET graft were fully characterized in vitro. This allowed us to compute the aortic pulse wave velocity (PWV) as *PWA* = *Z*_*C*_*A*/*ρ*, where *A* is the lumen cross-sectional area and *ρ* is the density of blood. Finally, we derived the aortic area compliance *C*_*A*_ from the Bramwell-Hill equation as 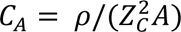, and the aortic distensibility *D* from its definition, as *D* = *C*_*A*_/*A*. The total arterial compliance and PWV are often measured in a clinical setting and they are preferred by physicians over the direct estimation of the local aortic compliance. They are associated with the arterial stiffness and the pulsatile arterial load that the heart encounters during ejection, while they are used as powerful biomarkers and predictors of cardiovascular risk ^33,34^.

### 2.7 Statistical analysis

Means were compared with repeated measures ANOVA. Multiple comparisons were corrected with the Fisher’s LSD procedure, which was proven to be the most powerful among alternative methods to correct multiple hypotheses tests, while still controlling the familywise error rate in the case of comparisons among three groups ^35^. Normality was determined by the Shapiro–Wilk test. The level of statistical significance for all analyses was set to 0.05. Values are reported as mean (standard deviation, SD). In all figures, the boxes represent the interquartile range, while the empty square and the whiskers represent the mean and the mean ± 1 SD, respectively.

## 3. Results

### 3.1 Surgery

The CM grafts could be handled and implanted using the same technique as standard grafts, without any surgical complications. All animals survived the operation, with the exception of one, whose descending aorta ruptured inferiorly of the distal anastomosis before the final measurement, because of microdamage to the arterial wall during the surgery. The rupture was not related to the implanted graft and it could not be sutured, due to its location. No other complications occurred during the surgical procedure.

### 3.2 Arterial waveforms

The replacement of the aorta with the CM graft did not significantly alter the shape of the pressure wave and increased only slightly the pulse pressure amplitude (Fig. 4). When only the gold-standard PET graft remained implanted, the pulse pressure amplitude increased, the aortic Windkessel function declined, while the pressure waveform was characterized by a late systolic increase and a prominent anacrotic inflection point in early systole. The aortic replacement with any of the two grafts did not result in significant differences in the aortic flow rate amplitude and wave shape.

**Figure 4.**
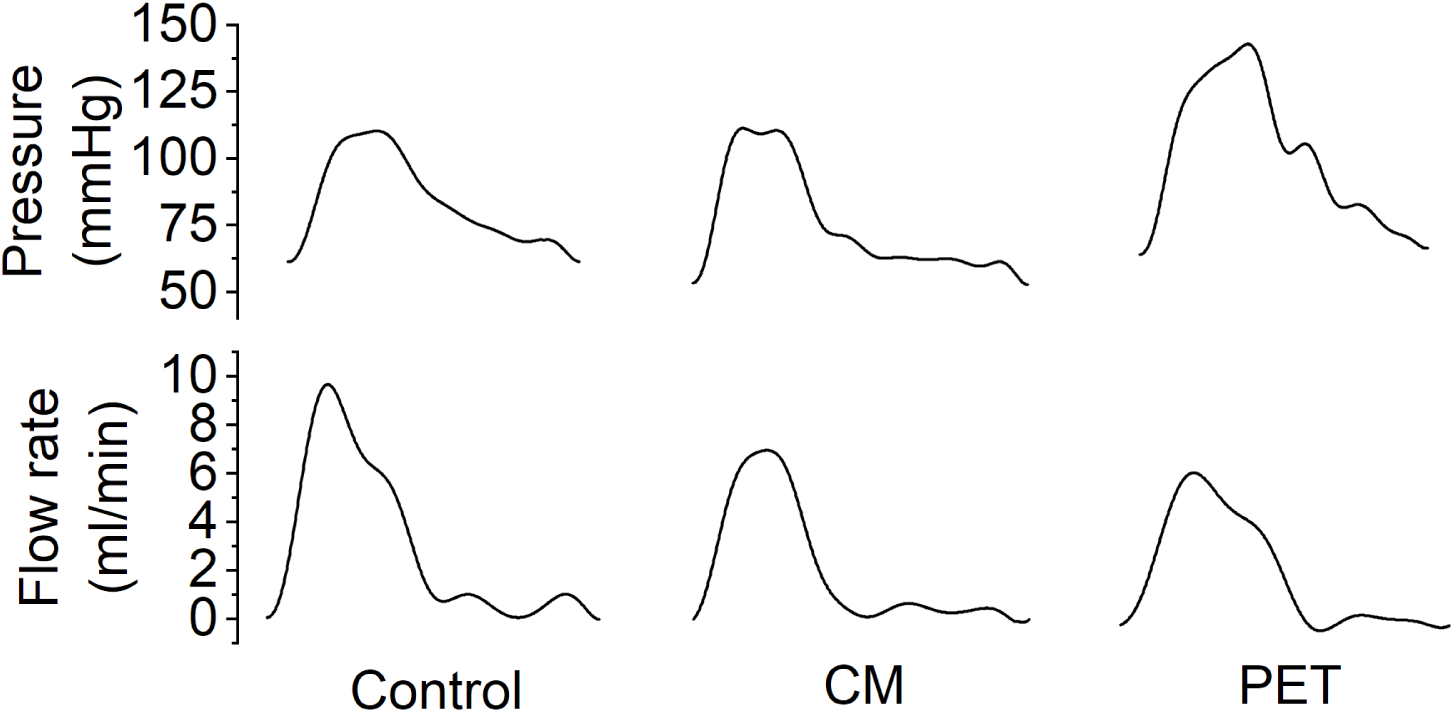
Representative aortic pressure and flow rate waveforms recorded in one animal. Compared to the initial measurement with the healthy aorta of the animal (Control), there was no significant difference in systolic pressure when the aorta was replaced with the compliance-matching graft (CM), while there was a small increase in pulse pressure. When only the gold-standard graft remained in place (PET), both the systolic and the pulse pressure increased, and a late systolic increase was observed in the pressure wave. The flow rate was not significantly altered in any of these cases.

### 3.3 Cardiovascular parameters

The measured or derived hemodynamic and arterial parameters are summarized in Table 1, along with the comparisons among the three conditions. The CM graft did not change the systolic pressure (*p* = 0.26), although it slightly but significantly (*p* = 0.018) increased the pulse pressure from 34.8 mmHg (*SD =* 8.3) to 56.9 mmHg (*SD =* 7.1). On the contrary, the PET graft significantly increased the systolic pressure from 90.5 mmHg (*SD =* 15.7) to 133.3 mmHg (*SD =* 13.3) compared to the healthy aorta (*p* = 0.005) and to CM graft (*p* = 0.031). Moreover, the PET graft augmented the pulse pressure more than twice the control value, that is 78.6 mmHg (*SD* = 15.7), which was significantly higher than the control (*p* < 0.001) and the CM (*p* = 0.018) cases.

**Table 1.** Hemodynamic and cardiovascular parameters, and the pairwise comparisons among the three conditions: healthy aorta (Control, n=6), compliance-matching graft (CM, n=6), and gold-standard graft (PET, n=5). Values are reported as mean (SD), * p < 0.05, ** p < 0.01, *** p< 0.001.

|  | Control | CM | PET | Con/CM | Con/PET | CM/PET |
| --- | --- | --- | --- | --- | --- | --- |
| Heart rate (min<sup>-1</sup>) | 107 (11) | 109 (15) | 121 (21) |  |  |  |
| Cardiac output (l/min) | 3.3 (0.6) | 2.7 (0.5) | 3.3 (0.5) |  |  |  |
| Stroke volume (ml) | 31.7 (7.5) | 24.8 (4.1) | 28.2 (7.9) |  |  |  |
| Maximum flow rate (l/min) | 11.3 (1.1) | 9.6 (4.0) | 12.5 (4.8) |  |  |  |
| Systolic pressure (mmHg) | 90.5 (15.7) | 100.5 (17.5) | 133.3 (13.3) |  | ** | * |
| Diastolic pressure (mmHg) | 55.7 (11.2) | 43.5 (17.2) | 54.7 (9.3) |  |  |  |
| Mean pressure (mmHg) | 67.3 (12.3) | 62.5 (16.9) | 80.9 (7.9) |  |  |  |
| Pulse pressure (mmHg) | 34.8 (8.3) | 56.9 (7.1) | 78.6 (15.7) | * | *** | * |
| Peripheral resistance (s mmHg/ml) | 1.22 (0.16) | 1.39 (0.25) | 1.53 (0.38) |  |  |  |
| Total arterial compliance (ml/mmHg) | 0.51 (0.25) | 0.20 (0.12) | 0.16 (0.09) | ** | ** |  |
| Characteristic impedance (s mmHg/ml) | 0.24 (0.09) | 0.33 (0.09) | 0.52 (0.16) |  | ** |  |
| Aortic area compliance (mm<sup>2</sup>/mmHg) | 0.62 (0.44) | 0.48 (0.19) | 0.17 (0.11) |  | * |  |

The CM graft managed to maintain the healthy levels of aortic characteristic impedance (*p* = 0.13) and aortic area compliance (*p* = 0.22), while the PET graft caused a significant rise (*p* = 0.004) in the characteristic impedance from 0.24 s mmHg/ml (*SD* = 0.09) to 0.52 s mmHg/ml (*SD* = 0.16), which resulted in a significant reduction in aortic area compliance (*p* = 0.03). Similar behavior was observed in the other arterial stiffness indices. The CM graft reduced PWV from 9.7 m/s (*SD* = 3.0) to 8.1 m/s (*SD* = 2.2), but this reduction was not statistically significant (*p* = 0.89). The PET graft, though, increased PWV to 16.0 m/s (*SD* = 5.0), a value that was significantly higher than both the PWV of the native aorta (*p* = 0.012) and the one of the CM graft (*p* = 0.009). The healthy aortic distensibility was 1.8 × 10^-3^ mmHg^-1^ (*SD* = 0.9 × 10^-3^) and it was maintained statistically the same (*p* = 0.67) with the CM graft, which had a distensibility of 2.5 × 10^-3^ mmHg^-1^ (*SD* = 1.0 × 10^-3^). The PET graft had a distensibility of 6.9 × 10^-4^ mmHg^-1^ (*SD* = 4.4 × 10^-4^), which was lower that the native aortic and CM graft distensibility (*p =* 0.047, and *p* = 0.024, respectively). The induced changes in systolic pressure, aortic PWV and distensibility can also be seen in Fig. 5.

**Figure 5.**
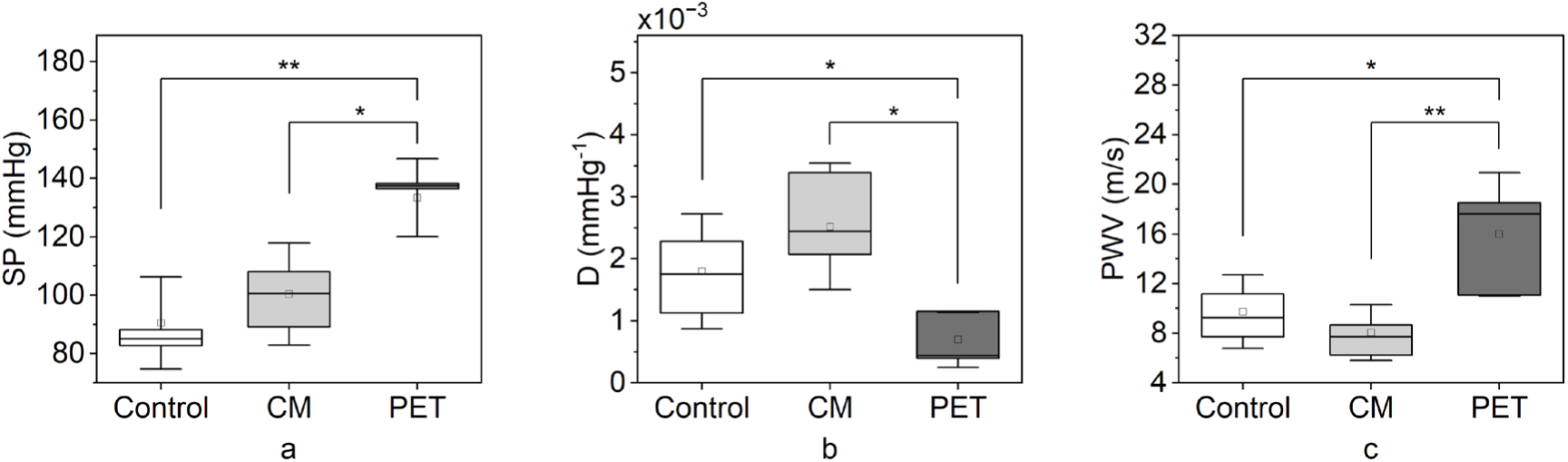
The compliance-matching graft did not significantly affect the systolic blood pressure and the aortic stiffness. (a) Systolic pressure (SP), (b) Aortic distensibility (D), and (c) Aortic pulse wave velocity (PWV) measured in the three different conditions. Control: healthy aorta, CM: compliance-matching graft, PET: PET graft, * p < 0.05, ** p < 0.01, non-significant differences are not designated.

The remaining cardiovascular parameters, such as the heart rate, the cardiac output, and the total peripheral resistance, did not change significantly due to the aortic replacement with both grafts (Table 1). Although the mean pressure and heart rate were elevated, potentially due to the reduction in compliance induced by the PET graft, the change was not statistically significant. These findings allowed a more appropriate reference for the comparison of the remaining parameters, that could be compared under similar hemodynamic baseline conditions. This was particularly important for pressure or heart rate dependent parameters, including PWV and distensibility. The only affected systemic parameter was the total arterial compliance, which was lowered after the replacement with the CM and PET grafts (*p* = 0.008 and *p* = 0.004, respectively).

## 4. Discussion

### 4.1 Main advantages of compliance-matching grafts

The mismatch of elastomechanical properties between the native artery and the vascular graft affects grafts of all sizes. Compliance mismatch impairs the performance of vascular grafts, which, currently, manifest suboptimal clinical outcomes with often deleterious consequences for the patients. There is a noteworthy absence of compliant grafts, especially large-diameter ones, while it remains unclear whether grafts with improved compliance could alleviate the complications caused by existing prostheses. Identification and verification of methods to enhance the agreement of elastic properties between arteries and grafts will facilitate the development of novel prostheses and may improve their clinical performance and the overall quality of treatment. In this study, we evaluated in a swine model the acute cardiovascular effects induced by recently proposed compliance-matching grafts, and compared these grafts to healthy, native aortas and to gold-standard, PET grafts.

Through these experiments, we demonstrated that the compliance-matching grafts could mimic the compliance of healthy aortas in vivo, and they could diminish the post-operative hypertension caused by the existing grafts. Thus, the proposed design led to the first graft to fulfill the, thus far, unmet need for compliant large-diameter grafts. The compliant graft diminished hypertension in a passive, purely biomechanical way, without any active components or bioactive compounds. Our findings provide further proof that post-operative hypertension, and potentially the associated complications of aortic grafts, are caused by the mismatch of mechanical properties and not by other underlying mechanisms.

### 4.2 Differences with existing approaches

The compliant grafts could maintain their mechanical properties in vivo, while their remaining characteristics, implantation procedure, and handling were similar to existing gold-standard prostheses. A previous study proposed bilayer stent-grafts that can achieve compliance close to the physiological aortic range, by adjusting the materials and the structure of the knitted part of the prostheses ^21^. However, the researchers used the stents only for adding mechanical support, resulting in lower-than-desired compliance. These grafts were not evaluated in vivo, while they could lose a major part of their compliance if they are extended longitudinally due to the periodic retraction of the aortic annulus ^36^ or if blood infiltrates and, subsequently, coagulates between the graft’s layers. Similarly, recently proposed, small-diameter, graft designs ^22–25^, even though they have been assessed in vivo, were tuned to match the compliance of peripheral arteries and not the inherently higher compliance of the aorta. It is also unclear how they would respond to the markedly different hemodynamic conditions and mechanical loads that prevail in the aorta. The CM graft design was demonstrated here as a large-diameter aortic graft, however, the same design approach with appropriate dimensions could be utilized to create compliant small-diameter grafts.

In contrast to the compliance-matching grafts, the gold-standard PET grafts had notably lower compliance than the healthy aorta (73% lower), as can be also seen from the PWV and distensibility comparison, and they caused an immediate rise in systolic pressure by 47%. The rise in pressure could be further amplified if the effect of anesthesia ceased, as sevoflurane causes vasodilation and reduces blood pressure ^37^. The increase in systolic pressure is the major shortcoming of existing, polymer-based, woven or knitted prostheses, and the chronically increased afterload leads to cardiac remodeling and hypertrophy ^17^. Our findings provide further proof for the suboptimal performance of existing prostheses. This is supported by the fact that arterial stiffness has been identified as an independent predictor of all-cause and cardiovascular mortality in humans, and other adverse outcomes, while an increase in PWV by one standard deviation increased cardiovascular events by 30% after adjustment for traditional risk factors ^38^. In our experiments, grafting with non-compliant prostheses increased PWV by 6.2 m/s or 64%, which corresponds to more than two standard deviations, and it is therefore associated with significantly increased risk for adverse outcomes. The measured systolic pressure with the PET grafts agrees with the reported systolic pressure ranges in similar experiments that involved either acute ^6^ or chronic ^39^ aortic bypass with inelastic grafts in dogs, and chronic aortic banding with PET bands in swine ^13,14^. In all those experiments, the pulse pressure approximately doubled after the bypass or the banding, which is in agreement with our current findings.

### 4.3 Aortic pressure waveforms

Interestingly, the examination of the pressure waveform revealed a prominent late systolic increase when only the PET graft remained implanted. Although the aortic pressure waveform types have not been classified in swine, the waveform in the healthy aorta and the CM graft resembled the human Type B pressure waveform ^40^. Type B waveform is characteristic of normotensive adults. We did not observe Type C waveforms, which are typical in younger individuals, despite the young age of the subjects. This could be attributed to the shorter length of porcine aortas compared to human ones, which does not allow enough time for consequent cardiac beats to overlap and create a Type C wave. The pressure waveform in the PET grafts resembled the Type A waveform, which is found in older, hypertensive adults and is characterized by increased pressure augmentation index, backward reflections and cardiac afterload.

### 4.4 Impact of the surgical and experimental procedure

The comparisons among the healthy aorta, the CM graft and the PET graft were performed at statistically equal conditions. The reference hemodynamic parameters, namely cardiac output, mean aortic pressure and total peripheral resistance, did not change among the three measurements. This allowed a more objective comparison. The only systemic parameter that was affected by the aortic replacement was total arterial compliance. The reduction in total arterial compliance could be attributed to the fact that we chose not to anastomose the supra-aortic trunk on the graft, to reduce the complexity of the surgery, as explained previously. Therefore, the cerebral and part of the upper body circulation were not included in the calculation of the total arterial compliance after the aortic replacement. Furthermore, even though we performed a cadaveric study to determine the size, both the CM and PET graft had smaller diameter than the native aortas, 18%–35% and 8%–27% respectively, resulting in reduced aortic volume compliance (23% and 73% lower respectively), which is a major part of the total arterial compliance. This fact could also explain why the CM graft caused a slight rise in systolic (11%) and pulse pressure (63%), even though such a behavior would not be expected from a graft with matching compliance. The size discrepancy between the aortas and the grafts was the main reason behind the selection of size-independent quantities, such as PWV and distensibility, as measures of aortic stiffness.

### 4.5 Limitations

The main limitation of our approach was that we performed acute animal experiments, instead of chronic ones. We selected the acute in vivo experiments as a necessary preliminary evaluation of our graft, given the increased complexity of the surgery. Nevertheless, similar experiments with stiff aortic banding in swine showed that there were only minor differences (less than 10%) in compliance, systolic, and pulse pressure between the recorded values directly after the surgery, after two days ^14^, and after two months, even when cardiac remodeling occurred ^13^. According to these results, the adverse cardiovascular effects caused by the non-compliant grafts, which we assessed in this study acutely, could remain nearly unchanged over time. The same could apply to the positive effects of matching compliance, as long as the graft could retain its compliance over time. Although the compliance of explanted grafts requires further investigation, it was suggested that woven and knitted polymer grafts gradually lose their compliance due to tissue ingrowth ^41,42^ or due to irreversible yarn rearrangement caused by cyclic loading ^43^. We expect that these two mechanisms will only moderately impact our CM graft, because the compliance of the CM graft is regulated by the stent layer in a manner nearly independent of the properties of the inner graft. The Nitinol stent is less prone to changes of mechanical properties due to the aforementioned mechanisms compared to polymer grafts.

Other limitations relate to the conditions under which the measurements were acquired. General anesthesia is known to influence the heart rate and blood pressure ^37^. Some administered drugs could also have cardiovascular effects, and the instrumentation could influence the results. We controlled for these effects by recording the reference hemodynamic conditions between measurements. The reference conditions were not statistically different between measurements, while the instrumentation remained the same throughout the experiments. As a result of these effects, the absolute values of the reported quantities might differ from the expected values under normal conditions. For example, the PWV values seemed slightly elevated compared to the expected values of healthy aortas. In spite of this fact, the conditions were similar between measurements and we focused on the comparison among the three cases and not on the absolute values per se. Finally, with these experiments we demonstrated that there is clinical significance in the use of the CM graft in terms of hemodynamic conditions, but this could not necessarily translate to improved clinical outcomes after chronic implantation. Further in vivo evaluation of the CM graft or other compliant grafts would be required to prove the long-term benefits of this type of grafts comparatively to existing alternatives.

## 5. Conclusion

In conclusion, we demonstrated by acute, in vivo experiments that the proposed compliance-matching graft can reach the compliance of a healthy aorta and address the absence of large-diameter, compliant, vascular prostheses. Importantly, the compliant grafts can surpass the clinical performance of gold-standard grafts, by mitigating the phenomenon of compliance mismatch and by alleviating post-operative complications, including graft-induced hypertension. These benefits were achieved by utilizing a multi-layer stent graft design that can be handled identically to existing prostheses, and it could be more resistant to the alteration of its mechanical properties after chronic implantation. Further research would be necessary to verify if compliance-matching grafts could lead to improved clinical outcomes. We believe that our design approach could pave the way for patient-specific compliant grafts of any diameter, while the proposed compliant grafts could become a noteworthy advancement towards the amelioration of the life expectancy and the quality of life of the patients that will receive them.

## 6. Acknowledgments

The authors would like to thank F. Fontao and X. Belin for their invaluable assistance in organizing and conducting the animal experiments, along with M. Malki and N. Levan for performing the cardiovascular perfusion.

## Notes

### Competing Interest Statement

The authors have declared no competing interest.

